# A potential role for epigenetic processes in the acclimation response to elevated *p*CO_2_ in the model diatom *Phaeodactylum tricornutum*

**DOI:** 10.1101/364604

**Authors:** Ruiping Huang, Jiancheng Ding, Kunshan Gao, Maria Helena Cruz de Carvalho, Leila Tirichine, Chris Bowler, Xin Lin

## Abstract

Understanding of the molecular responses underpinning diatom responses to ocean acidification is fundamental for predicting how important primary producers will be shaped by the continuous rise in atmospheric CO_2_. In this study, we have analyzed global transcriptomic changes of the model diatom *Phaeodactylum tricornutum* following growth for 15 generations in elevated *p*CO_2_ by strand-specific RNA sequencing (ssRNA-seq). Our results indicate that no significant effects of elevated *p*CO_2_ and associated carbonate chemistry changes on the physiological performance of the cells were observed after 15 generations whereas the expression of genes encoding histones and other genes involved in chromatin structure were significantly down-regulated, while the expression of transposable elements (TEs) and genes encoding histone acetylation enzymes were significantly up-regulated. Furthermore, we identified a series of long non-protein coding RNAs (lncRNAs) specifically responsive to elevated *p*CO_2_, suggesting putative regulatory roles for these largely uncharacterized genome components. Taken together, our integrative analyses reveal that epigenetic elements such as TEs, histone modifications and lncRNAs may have important roles in the acclimation of diatoms to elevated *p*CO_2_ over short time scales and thus may influence longer term adaptive processes in response to progressive ocean acidification.

## Introduction

Ocean acidification is caused by the absorption of excessive anthropogenic CO_2_ emissions by the ocean. It is predicted that the further rise of atmospheric CO_2_ concentrations under a “business-as-usual” CO_2_ emission scenario will lead to a further drop of ocean surface pH by 0.4 units (1). Diatoms, responsible for about 20% of global primary production, profoundly influence marine food webs and biogeochemical cycles (2). Although several studies reporting the effects of ocean acidification on diatoms at the physiological and ecological levels have been reported (3), there are only limited studies deciphering the underlying molecular mechanisms of diatoms in response to ocean acidification.

Most studies that have examined the genetic responses of diatoms to ocean acidification have focused on carbon fixation and assimilation genes. It has been reported that the large subunit of ribulose-1, 5-bisphosphate carboxylase/oxygenase (RubisCO), involved in the first committed step of carbon fixation, was down-regulated at the transcriptional and protein levels under elevated *p*CO_2_ conditions (4, 5). Genes encoding components of carbon concentrating mechanisms (CCMs) including carbonic anhydrase and bicarbonate transporters, which concentrate CO_2_ against the cellular gradients and promote delivery of CO_2_ to the RubisCO active site as a means to outcompete the non-productive oxygen fixation reaction, were also reported to be down-regulated under elevated CO_2_ conditions (6–8). Furthermore, using RNA-seq in the context of a nitrate-limited chemostat experiment, it was additionally reported that elevated CO_2_ first caused changes in transcriptional regulation and chromosome folding and subsequently metabolic rearrangements in *Thalassiosira pseudonana.* This study further revealed that genes in one CO_2_-responsive cluster share a putative cAMP responsive *cis*-regulatory sequence for a cAMP second messenger system (9). These results imply that in addition to genes encoding central components of carbon fixation and assimilation, the response to elevated CO_2_ may involve a broader reprogramming of transcription, genome structure, and cell signaling.

Long non-protein coding RNAs (lncRNAs) are transcripts longer than 200 nucleotides which are similar to mRNAs but have low to no apparent protein coding potential. Long considered as transcriptional noise, numerous evidence from animals and plants gathered over recent years indicate that lncRNAs have multiple and important functions, such as histone modification regulation, transcription machinery regulation and post-translational regulation (10, 11) Furthermore, lncRNAs are often associated with stress responses having fundamental regulatory functions in cellular adaptation and homeostasis (12). LncRNAs may also have important roles in diatoms in response to different environmental stimuli, however to our knowledge only one study has so far investigated changes in lncRNA levels in response to phosphate fluctuations (13).

Histone play crucial roles in genome stability. Two copies of the core histones (H2A, H2B, H3, and H4) are wrapped with ca. 147 bp DNA to form the nucleosome core particle. Up to 80 bp of linker DNA with linker histone H1 connects nucleosome core particles along the DNA (21). A total of 14 histone genes (H2A, H2B, H3, H4, H1) are dispersed on 5 of the 34 chromosome scaffolds of the *P. tricornutum* genome (15, 22). Transposable elements (TEs) are mobile DNA sequences able to move in the genome by generation of new copies. TEs are ancient and active genomic components which have been found in most eukaryotic and prokaryotic genomes (14). In *P. tricornutum* and *T. pseudonana*, 5.8% and 1.1% of the genome are contributed by TEs, respectively (15–17). TEs are classified into retrotransposons (Class I) and DNA transposons (Class II) according to the presence or absence of an RNA transposition. Class I transposons use a “copy and paste” strategy to proliferate in the genome whereas the majority of Class II transposons use a “cut and paste” strategy to move in the genome. Class I retrotransposons include Long Terminal Repeat (LTR) Elements and non-LTR elements (such as LINE-type TEs). A total of 90% and 58% of TEs are LTR retrotransposons (LTR-RTs) in the *P. tricornutum* and *T. pseudonana* genomes, respectively. Among LTR retrotransposons, *gal pol* domains are the most abundant TEs in *P. tricorntutum* (17, 18). Retrotransposons are considered as the major contributors to the expansion of genomes especially in higher eukaryotes and enhance their hosts’ fitness by advantageous genome re-arrangements in a changing environment (19, 20). In the current study, we have combined physiological performance measurements, with strand-specific RNA sequencing (ssRNA-Seq) to assess global transcriptomic changes in *P. tricornutum* in response to elevated *p*CO_2_ with respect to ambient *p*CO_2_ concentrations following growth for 15 generations. Our data provide new insights into how a diatom copes with elevated *p*CO_2_ using some potential epigenetic mechanisms and suggest an involvement of TEs and possibly lncRNAs in the acclimation response to elevated *p*CO_2_.

## Materials and methods

### *P. tricornutum* culture conditions and physiological measurements

Axenic cultures of *Phaeodactylum tricornutum* (Bohlin) strain CCMP 632 were obtained from the Culture Center of Marine Phytoplankton (East Boothbay, ME, USA). *P. tricornutum* cells were maintained in polycarbonate bottles (Nalgene, USA) at 20 °C under cool white fluorescent lights at 150 μmol m^2-^ s^-1^ (12h:12h light: dark). Different *p*CO_2_ in the medium was achieved by bubbling different concentrations of CO_2_. The gas streams were pre-filtered through 0.2-μm HEPA filters. The low CO_2_ medium was bubbled with ambient air of about 400 ppmv (low CO_2_, LC) and the high CO_2_ medium was bubbled with pre-mixed air-CO_2_ mixtures (1000 ppmv; high CO_2_, HC) from a plant growth CO_2_ chamber (HP1000G-D, Ruihua) at a flow rate of 0.6 L min^−1^, controlling the CO_2_ concentration with a less than 3% variation. Triplicates of LC and HC cultures were all started with the same initial cell densities of 2 × 10^4^ cells mL^−1^. The culture medium was refreshed with *p*CO_2_ and the pH adjusted every 48 hours to maintain cell concentrations ranging from 2 × 10^4^ to 2 × 10^5^ cells mL^−1^ and thus maintaining the desired pH and *p*CO_2_ values within a variation of less than 0.5%. The carbonate system in the HC cultures differed significantly from the LC culture (Supplementary Table 1). The growth rate was calculated as μ = (ln C_1_–ln C_0_)/(t_1_–t_0_), where C_0_ and C_1_ represent the cell density at t_0_ (initial or just after the dilution) and t_1_ (before the dilution), respectively. Cell densities were measured by particle count and size analyzer (Z2 Coulter, Beckman). Pigment concentrations were measured as described in (23) using spectrophotometry (Beckman DU-800, USA). Fv/Fm, effective quantum yield (PSII) were measured by a Xenon-Pulse Amplitude Modulated Fluorometer (XE-PAM, Walz, Germany) at the middle of the light period. Particulate organic nitrogen (PON) and carbon (POC) were measured according to (24).

### RNA extraction, library preparation and ssRNA-seq

*P. tricornutum* cells from three biological replicates were harvested in the middle of light period by vacuum filtration on 2 μm polycarbonate filters (Millipore, USA) following 15 generations (day 10), flash frozen in liquid N_2_ and maintained at −80 °C until use. Total RNA was extracted from frozen cell pellets using TRIzol^®^ Reagent (Invitrogen, Carlsbad, CA, USA) according to the manufacturer’s instructions. RNA integrity was assessed using the RNA Nano 6000 Assay Kit of the 2100 Bioanalyzer (Agilent Technologies, CA, USA) and by RT-qPCR. A total amount of 3 μg RNA per sample was used for construction of RNA sample libraries. Ribosomal RNA was removed by Epicentre Ribo-Zero™ Gold Kits (Epicenetre, USA). The sequencing libraries for ssRNA-Seq were generated using NEBNext Ultra Directional RNA Library Prep Kit (NEB, Ispawich, USA) with varied index labeling following the manufacturer’s guidelines. After cluster generation by HiSeq PE Cluster Kit v4-cBot-HS (Illumina), the libraries were sequenced on an Illumina Hiseq 4000 platform and 150 bp paired-end reads were generated. Library preparation and ssRNA-seq were performed offsite at Annoroad Gene Technology Corporation (Beijing, PR China).

### ssRNA-seq data assembly and analysis

Raw data from ssRNA-seq were cleaned up by removing adapter-contaminated reads (more than 5 adapter-contaminated bases), poly-N containing (more than 5%) and low quality reads (quality value less than 19, accounting for more than 15% of total bases). All the downstream analyses were based on filtered clean data with high quality. The average clean data was 7.1 Gb for ssRNA-Seq. The reference genome of *P. tricornutum* was downloaded from (http://protists.ensembl.org/Phaeodactylum_tricornutum/Info/Index). Bowtie2 v2.2.3 was used for building the genome index (25), and clean data was mapped to the reference genome using TopHat v2.0.12 (26). Read counts for each gene in each sample were counted by HTSeq v0.6.0, and RPKM (Reads Per Kilobase Millon Mapped Reads) was then calculated to estimate the expression level of genes in each sample (27).

DESeq (v1.16) was used for differential gene expression analysis. The *p*-value was assigned to each gene and adjusted by the Benjamini and Hochberg’s approach for controlling the false discovery rate. Genes with fold change ratio (HC/LC) ≥2 (q≤0.05) and fold change ratio ≤0.5 (q≤ 0.05) were defined as “up-regulated genes (UGs)” and “down-regulated genes (DGs)”, respectively. Genes with fold change ratio (HC/LC) between 1.5 and 2 (q≤ 0.05), fold change between 0.5 and 0.67 (q≤ 0.05) were defined as “relatively up regulated genes (RUGs)” and “relatively down regulated genes (RDGs)”, respectively.

### GO and KEGG analysis

Gene Ontology analysis (http://www.geneontology.org) was conducted to construct meaningful annotation of genes and gene products in different databases and different species. GO enrichment analysis was used to assess the significance of GO term enrichment among differentially expressed genes (DEGs) in the domains of biological processes, cellular components and molecular functions. The GO enrichment of DEGs was implemented by the hypergeometric test, in which *p*-value was calculated and adjusted as *q*-value. GO terms with *q*<0.05 were considered to be significantly enriched. We further performed KEGG (Kyoto Encyclopedia of Genes and Genomes, http://www.kegg.jp/) pathway analysis to explore the roles of the differentially expressed genes in different metabolic pathways and molecular interactions. The genes related to chromatin, protein DNA, chromosomal part, chromosome, histone modification, glycolysis TCA cycle, and photosynthesis (light part), carbon assimilation and N uptake and assimilation were classified manually based on the KEGG and GO gene classification (Supplementary Table 2).

### Identification of novel IncRNAs

LncRNA candidates with length of ≥ 200 nucleotides, a predicted open reading frame (ORF) of ≤ 100 amino acids, number of exon ≥ 1 and reads ≥ 3 were selected (13, 28). Transdecoder was used to remove known mRNA transcripts and candidates with protein coding potential. The novel lncRNAs were classified based on the location of the lncRNA gene region in the reference genome. In order to increase the depth of the reads for the increased detection of noncoding RNA transcripts, the noncoding RNA transcript reads derived from the three replicates were pooled since the Pearson correlation coefficients (PCCs) between the RPKM of three replicates of LC and HCs were high (> 0.97). A 1.5-fold /0.67-fold (up-regulated/down regulated) and variance in RPKM and a *p* value (Fisher’s exact test) of < 0.05 were used as cutoffs to define differentially expressed genes, respectively.

### Interaction network construction

The *P. tricornutum* protein–protein interaction (PPI) network data was downloaded from http://string-db.org/. The differentially expressed mRNA interaction networks, specifically transcription factor (TF), histone modification genes and mRNAs interaction networks were constructed in Cytoscape 3.4.0 based on the *P. tricornutum* PPI network data database.

### Quantitative real-time reverse transcription PCR

Reverse transcription quantitative PCR (RT-qPCR) was used to verify the transcriptomic data. The primers used for qPCR are listed in Supplementary Table 3. Polyadenylated RNA was converted to cDNA by PrimeScript™ II Reverse Transcriptase (Takara). The RT product was then used as the template for qPCR. All qPCR reactions were performed on a FTC2000 (Canada) using SYBR Green I mix (Takara) in 96-well plates according to the manufacturer’s recommendations. TBP (TATA box binding protein) and RPS (ribosomal protein small subunit 30S), two housekeeping genes (29), were used as references to calibrate the expression. Three technical and three biological replicates were conducted.

## Results and Discussion

### Physiological response of *P. tricornutum* to elevated *p*CO_2_

No significant differences were detected between cells grown under the HC and LC conditions with respect to pigment contents, Fv/Fm, photosynthetic yield, POC and PON content after growth for 15 generations (Table 1). However, a slight increase in specific growth rate (2.6%) was observed in the HC condition, which is consistent with our previous study (30). The lack of physiological differences between LC and HC after 15 generations suggests an effective and stable acclimated state to the new CO_2_ saturated environment, revealing a very dynamic plasticity to environmental change in diatoms.

### The global transcriptomic response to elevated *p*CO2 after growing for 15 generations

Strand specific (ss)RNA-seq data generated from the HC and LC physiological states were assembled and mapped. The average mapping rate for ssRNA-seq data was about 76.8%. The mapping rate of reads is shown in Supplementary Table 4. The average mapping rate on the intergenic regions was 24.8% as shown in Supplementary Table 4. The expression profile of each gene is listed in Supplementary Table 5. The GO enrichment analysis of DEGs in the domains of biological processes, cellular components and molecular functions is shown in Supplementary Figure 1. In general, the numbers of differentially regulated genes under the HC condition were much lower than induced by other stimuli such as nitrogen starvation and phosphate depletion (13, 31). This may suggest that elevated *p*CO_2_ is not a strong environmental stimulus compared to nitrogen starvation and phosphate depletion for *P. tricornutum.* Globally, 600 protein coding genes were differentially expressed in response to HC with 191 UGs, 287 RUGs, 175 DGs and 313 RDGs compared with the LC condition over 15 generations. A good correlation between transcript abundance was found between transcriptomic data and RT-qPCR, thus validating the robustness of the sequencing data (Supplementary Table 6).

### The impact of elevated *p*CO_2_ on metabolic genes

Our results showed that some genes involved in CCM, energy-production/consumption and light capture were down-regulated after growing in the HC condition for 15 generations (Figure 1, Figure 5). As expected, some genes involved in CCM were down-regulated since high CO_2_ is expected to reduce the need for CCM. Two genes encoding pyrenoid-localized carbonic anhydrases (Pt CA1: Phatr3_J51305, Pt CA2: Phatr3_J45443) were down-regulated in the HC condition while other genes encoding carbonic anhydrases were not significantly affected by changing CO_2_ concentrations (32). Based on RT-qPCR results, the transcriptional level of the putative carbonic anhydrase Phatr3_J51305 showed significant down-regulation in HC compared to LC (Supplementary Table 6). The putative HCO_3_^-^ transporter (Phatr3_ J54405, SLC4-5) was also down-regulated in HC. However, other putative HCO_3_^-^ transporters, including SLC4-2 (Phatr3_Jdraft1806) which has been shown to encode a Na^+^-dependent HCO_3_^-^ transporter at the plasma membrane, were not significantly affected by HC in our study. Phatr3_J38227 encoding a putative CO_2_ hydration protein (ChpXY), which was not described in previous studies, was down regulated. The malate dehydrogenase gene, involved in the C4 pathway (Phatr3_J42398), showed repressed gene expression in the HC condition compared to the LC condition whereas other genes involved in C4 metabolism were not significantly affected. Furthermore, some genes encoding proteins with roles in energy-producing metabolic pathways, such as oxidative phosphorylation and TCA cycle, were preferentially down-regulated. These include genes encoding phosphoglycerate kinase (Phatr3_J48983), mitochondrial malate dehydrogenase (Phatr3_42398) and mitochondrial glyceraldehyde 3-phosphate dehydrogenase (Phatr3_25308). This suggests a general reduction in metabolism with an energy saving status in the HC acclimated state. Other genes encoding plastidial glyceraldehyde 3-phosphate dehydrogenases such as Phatr3_22122 and Phatr3_32747, albeit lacking clear subcellular localization information, were also down-regulated. Genes encoding components involved in electron transfer and oxidation/reduction such as NADH dehydrogenase (Phatr3_EG01423, Phatr3_EG00870) were preferentially down-regulated. Genes encoding V-type ATPase subunit F (Phatr3_J31133) and mitochondrial F-type ATPase (Phar3_J39529), which are directly related to energy consumption and production, respectively, showed down-regulation in HC. The down regulation of genes involved in CCM and energy turnover rate suggest that a general reduction in metabolism may have contributed to the slightly increased growth rate (2.6%) in HC conditions, which is also consistent with previous experimental and modeling results (6, 30).

**Figure 1.**
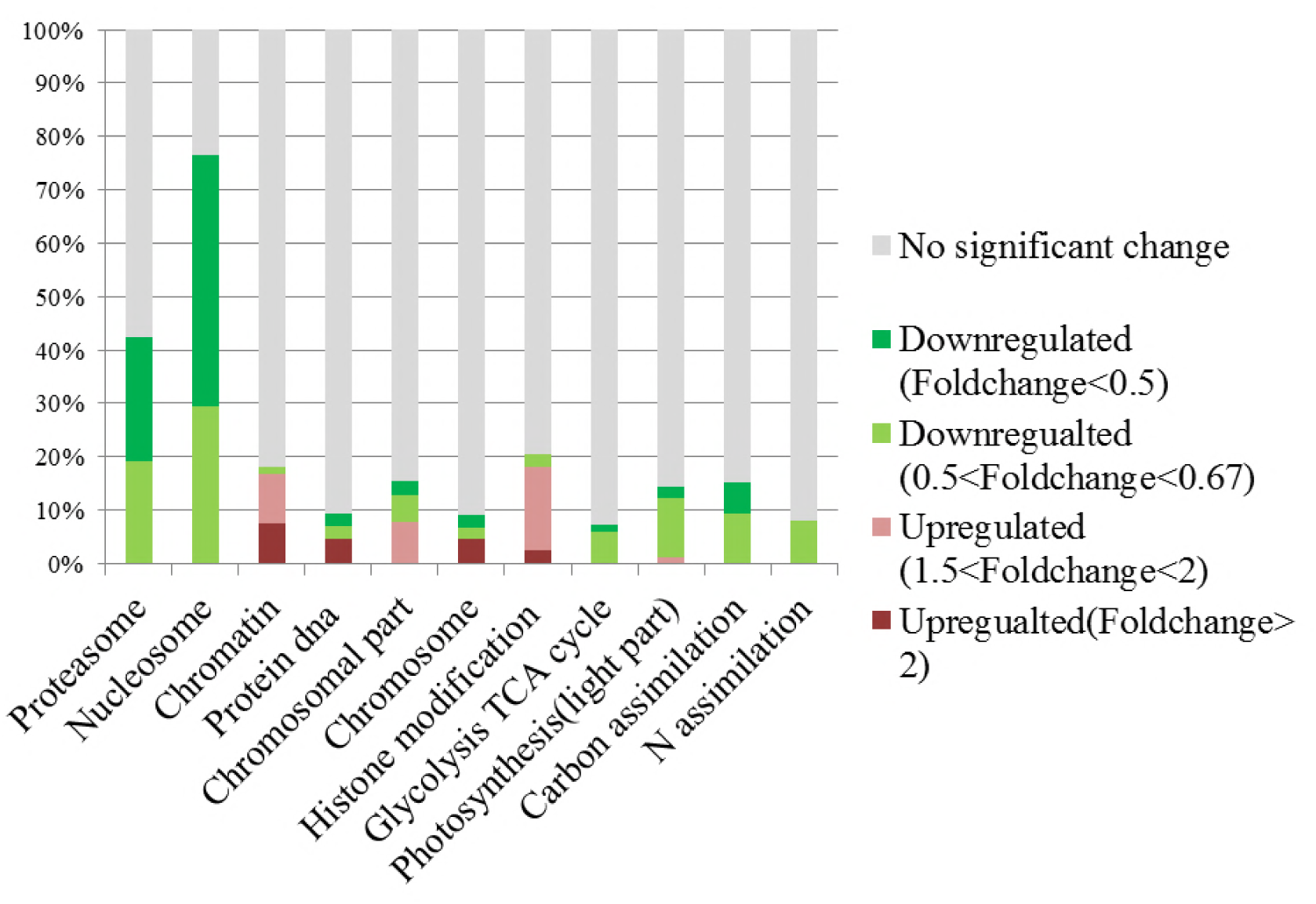
Transcripts abundance (HC/LC) of genes from 11 gene families after growing for 15 generations. Changes are denoted as the percentage of up-regulated with fold change⩾2 (dark red, UGs), up-regulated with 1.5⩾fold change>2 (light red, RUGs), down regulated with fold change⩽0.5(dark green, DGs), down-regulated with 0.67<fold change<0.5 (light green, RDGs), no significant changes (gray) within each family. The full description of the gene families and their fold change, false detection rate (FDR), and description can be found in Supplementary Table 4

Our physiological assays indicated that the photochemical performance was not significantly influenced after 15 generation under elevated *p*CO_2_; however, some light capture genes such as Fucoxanthin-chlorophyll a-c binding protein (Phatr3_J24119, Phatr3_J25893, Phatr3_J18049) and Fucoxanthin-chlorophyll a/c protein were down-regulated in the acclimated HC condition. RT-qPCR results also showed that the transcriptional level of Phatr3_J27278 was significantly down-regulated in HC (Supplementary Table 6). A gene encoding a putative Cyt b6f iron-sulfur subunit (Phatr3_J13358), which is essential for one of the light reactions in photosynthesis, was also down-regulated. Although photorespiration was not measured in our study, the putative 2-phosphoglycolate phosphatase gene (Phatr3_Jdraft1186), a key enzyme in the photorespiration pathway, was down-regulated indicating possible decrease in photorespiration rates under the HC condition, which is in line with the previous results in *T. pseudonana* (9).

A significant proportion of ribosome genes were down-regulated in HC while a significant proportion of genes encoding proteasome core complex were significantly down-regulated by elevated *p*CO_2_ (Figures 1, 5). The down-regulation of genes encoding proteins involved in proteasome function suggest that the protein degradation system was relatively inactive while the significant down-regulation of ribosome components may reflect either a decreased capacity for protein synthesis or a low turnover of ribosomal proteins in the acclimated HC condition. All these results indicate a low turnover rate of protein synthesis and degradation, which is somewhat in accordance with the lower energy cost implied by the down-regulation of energy producing genes in HC described above. The down-regulation of ribosome genes in the HC condition was in contrast with the previous study on *T. pseudonana* acclimated to high CO_2_ for 10 days (9), probably because the previous study involved low nitrate concentration. As reported in (33), Pt CA1 (Phatr3_J51305) was down-regulated under 5% CO_2_ (50000ppmv). Therefore, Pt CA1 (Phatr3_J51305) may be the most responsive carbonic anhydrase gene in face of elevated CO_2_ and essential in preventing the passive diffusion of concentrated CO_2_ out of the cell.

### The impact of elevated *p*CO_2_ on genes involved in pH homeostasis maintenance

Beside genes directly involved in metabolic pathways being affected by elevated *p*CO_2_, some genes related to cellular pH homeostasis maintenance were also affected by elevated *p*CO_2_. It is predicted that the bulk seawater H^+^ concentration 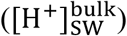 will increase by 100-150% by 2100 in response to ocean acidification (34). It was reported that the increased pH in seawater results in greater diel variation of H^+^ concentration proximate seawater 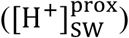 of marine organisms by boundary-layer processes (34). The change in 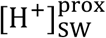 affects the operation of cellular processes involved in pH homeostasis including H^+^ production/consumption and active/passive H^+^ transport (35, 36). It has been reported that the down-regulation of CCM changes intracellular H^+^ concentration by decreasing HCO_3_^-^ transport into cytoplasmic matrix and conversion from HCO_3_^-^ to CO_2_ (36). To maintain the neutral and acid pH in cytoplasmic matrix and organelles, the proton pump plays critical roles in H^+^ transport (36). In our study, Phatr3_J31133 encoding the subunit F of vacuolar V-type proton pump where H^+^ is transported from cytoplasmic matrix into vacuoles coupled with ATP hydrolysis was down-regulated in HC; however, Phatr3_J43207 coding inorganic H^+^ pyrophosphatase was up-regulated stimulating H^+^ transport from cytoplasmic matrix into vacuoles coupled with pyrophosphate hydrolysis (Figure 5). Similarly, Phatr3_J39529 encoding epsilon chain of mitochondrial F-type proton pump in which H^+^ is transported from mitochondria into cytoplasmic matrix coupled with ATP synthesis was down-regulated; however, Phatr3_J50171 encoding protein related to mitochondria K^+^-H^+^ exchange was up-regulated (Figure 5). These results suggest that the decrease in H^+^ transport coupled with ATP metabolism may be offset by other H^+^ transport pathways to maintain the cellular pH homeostasis under high *p*CO_2_ conditions in *P. tricornutum.* More investigations, such as pH measurements in different organelles and the cytoplasm under high *p*CO_2_ conditions compared to ambient conditions, are needed for better understanding of pH homeostasis mechanisms operating under conditions reflecting ocean acidification in diatoms.

### Potential changes in epigenetic regulation in response to elevated *p*CO_2_ over 15 generations

After growing in HC and LC conditions for 15 generations, the most striking molecular response was the down-regulation of genes encoding histones and the up-regulation of many transposable elements (TEs). TE and histone gene variations and their changes in expression may have fundamental effects on the adaptive responses to elevated *p*CO_2_, as a result stabilizing the physiological performance. 12 out of 14 histone genes (H3, H4, H2B, H3B, H1) were significantly down-regulated. For example, Phatr3_J54360 encoding H2B 1b isoform showed down-regulation in both RT-qPCR (Supplementary Table 6) and ssRNA-seq data sets in HC. Only Phatr3_28445 encoding H2A 1 (H2A.Z) and Phatr3_J21239 encoding H3.2 were not significantly affected by elevated *p*CO_2_. Phatr3_EG01358 encoding histone H2A isoform 3a, with fold change 0.098 (HC/LC), was the most significantly down-regulated histone gene. The global down-regulation of genes encoding histones in response to HC may lead to chromatin structural changes and thus facilitate genome rearrangements.

On the other hand, genes encoding core domains of TEs, in particular the *gal pol* genes, were highly expressed in HC compared to LC condition. The TEs of the LTR-RTs are constituted by *gal pol* genes flanking with LTR (long terminal repeat) sequences. The *gal* gene encodes the virus-like particle (VLP) structural proteins where reverse transcription takes place while the *pol* genes encode several enzymes, the protease (PR) cleaving the POL polyprotein. LTR-RTs are classified into *Ty1/copia* elements and *Ty3/gypsy* elements based on the organization of their pol genes and similarities among their encoded RT proteins. Phatr3_EG00775, Phatr3_Jdraft1453 and Phatr3_J33646 encoding *gal pol* polyprotein belonging to *Copia* type elements were significantly up-regulated with fold changes of 4.73, 6.39 and 2.37 (HC/LC), respectively. Phatr3_Jdraft1453, encoding integrase core domain responsible for mediating integration of a DNA copy of the TE into the genome, also showed up-regulation in HC compared to LC. Phatr3_48048 encoding reverse transcriptase (RNA-dependent DNA polymerase) was up-regulated in HC compared to LC. The putative endonuclease genes (Phatr3_J44711, Phatr3_35971 Phatr3_J23306, Phatr3_J12645, Phatr3 J43711, Phatr3_J44711 Phatr3_J46452, Phatr3_EG00704, Phatr3_22703 and Phatr3_44042) and putative Ribonuclease H (Phar3_EG02045 and Phar3_J44863) integrating the cDNA into the genome were also up-regulated in HC compared to LC.

Among these active TEs, *Copia* type retrotransposons were the most active transposons in the HC condition compared to LC based on the order of core domain in the *pol* region. Based on the TE sequences provided in (18), CoDi 1.1 (EU432476.1, Phatr3_EG00052), CoDi 4.4 (EU432484.1, Phatr3_EG01511), CoDi 6.5 (EU432496.1, Phatr3_EG00775), CoDi 2.6 Blackbeard (EU363804.1, Phatr3_J50428) and CoDi 3.1 (EU432481.1, Phatr3_J33646) were significantly up-regulated in response to the HC condition, as shown in Figure 2. CoDi1.1 with fold change 73.82 (HC/LC) was the most responsive TE in response to HC. In *P. tricornutum*, it has been reported that Blackbeard and Surcouf, belonging to the *Ty1/copia* TEs, were responsive to nitrate starvation and high decadienal condition, respectively. In our study, the activation of *Copia* TEs, especially CoDi1, in response to the HC condition after growing for 15 generations was consistent with what has been observed in natural environmental samples that *Copia* TEs were more active compared to other TEs (37). This suggests that certain TEs specifically respond to given conditions and further emphasize the importance of *Copia* TEs in response to different environmental stimuli.

**Figure 2.**
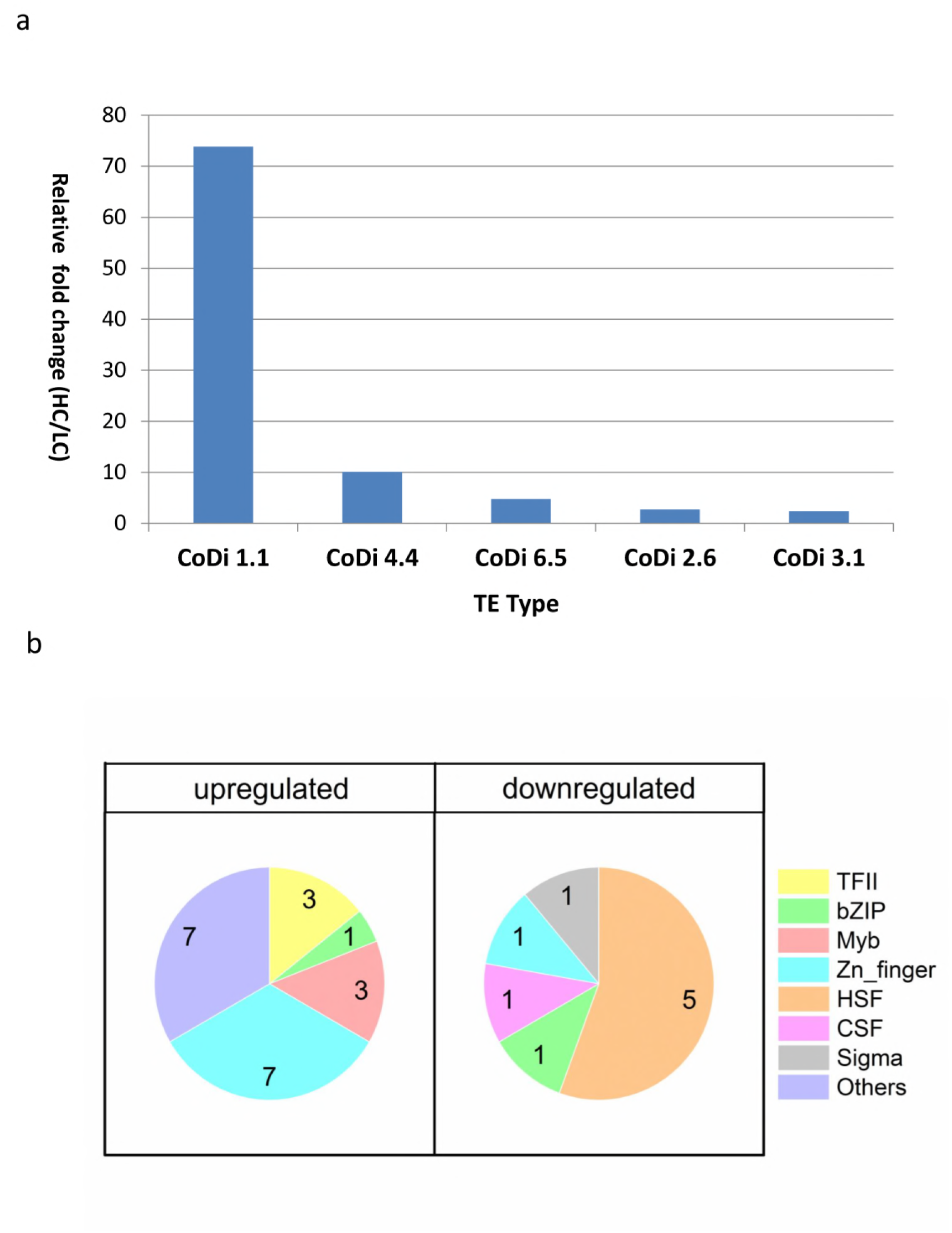
(a) The expression levels of different classes of transposon elements (HC/LC). (b) Pie charts of the different classes of up-regulated and down-regulated transcription factors (TFs) under the HC condition compared to the LC condition in *P. tricornutum* after growing for 15 generations (up-regulated: HC/LC fold change⩾1.5, down-regulated: HC/LC fold change⩽0.67, *P* < 0.05).

The mobile nature of TEs can contribute to alterations in genetic regulatory elements, therefore altering levels of gene expression, triggering genome rearrangements and mutations that can accelerate biological diversification and consequently influence genome evolution (38, 39). It is generally considered that the insertion of TEs has detrimental effects, including gene inactivation and alteration of chromosome structure. On the other hand, the beneficial effects of TEs have also been noted. TEs can enhance their hosts’ fitness, induce advantageous rearrangements or enrich the host’s gene pool in the long term (19, 20). It has been further recognized that TEs as diversifying agents may contribute to adaptive processes occurring over short time scales. These aspects may explain the abundance and ubiquity of TEs in the natural environment (38, 40, 41).

In our study, the activation of TEs accompanying the down-regulation of histone genes in the HC condition may increase the potential for genome restructuring, and consequently to reshape gene expression patterns by TE insertions in promoters, enhancers and exons, and result in sequence expansion, gene duplication, novel gene formation and expansion, and re-wiring of genetic regulatory networks. More importantly, the effects on the genomes induced by elevated *p*CO_2_ on the acclimated state could be transmissible to future generations, which not only increase the fitness and phenotypic variation for acclimation in short time but also tremendously influence the adaptation and evolution of the whole population in the long term under the elevated *p*CO_2_ conditions. Accumulating data has shown that TEs are not only the most abundant and ubiquitous genes in nature but also are transcriptionally active in marine assemblages and in mutualistic endosymbionts, which further indicate their importance in response to environmental change (37, 42, 43)

DNA methylation and histone modifications are two major epigenetic components, which can mediate heritable changes in gene functions beyond DNA sequence changes. In *P. tricornutum*, cytosine DNA methylation is commonly found in TEs and may be involved in controlling TE mobility in the genome (44). As a case in point, the activation of the retrotransposon *Blackbeard* was accompanied by its hypomethylation under nitrate starvation (18). The “epi-transposon” hypothesis proposes that changing environments can lead to stress-induced breakdown of epigenetic suppression of TEs (such as DNA methylation), which results in extensive transposition, thus providing new material for rapid adaptive shifts in short term acclimation (45). It was reported that some TEs lose DNA methylation in *P. tricornutum* in response to nitrate starvation (46). In our study, besides the down-regulation of histone genes, the burst of TEs was the most significant genetic response in the acclimated HC state. We therefore speculate that the highly expressed TEs may be accompanied by DNA demethylation on TEs in response to elevated *p*CO_2_, although there is currently a lack of genome wide DNA methylation data under the HC and LC conditions.

Previous studies have also shown that histone modification may play a role in responses to different environmental stimuli in diatoms. Histone acetylation is usually considered as the positive hallmark for gene expression and H3K9/14 acetylation and H3K4me2 marks were found to be modulated in response to nitrate limitation in *P. tricornutum* (46). It was further reported that genes encoding putative proteins involved in epigenetic modifications were up-regulated under phosphorus depletion in *P. tricornutum* (13). In the current study, the genes encoding proteins responsible for histone acetylation (Phatr3_J45764, Phatr3_J54343, Phatr3_J45703, Phatr3_EG02442, Phatr3_J45644) were up-regulated in HC. Further studies on whether the up-regulation of histone acetylases leads to acetylation of specific histones residues resulting in up-regulation of histone acetylation marked genes under elevated *p*CO_2_ conditions in *P. tricornutum* will therefore be of interest.

In order to further analyze the potential roles of histone modification in response to HC, the network of differentially expressed putative histone modification genes and their associated genes was constructed in Cytoscape (Figure 3). All putative histone modification genes, including putative histone deacetylase Phatr3_4423, putative histone acetyltransferases Phatr3_45703, Phatr3_54343 and Phatr3_45764 and putative H3K4 methyltransferase Phatr3_44935 were up-regulated in the constructed network in HC. The putative H3K4 methyltransferase (Phatr3_44935), histone deacetylase genes (Phatr3_4423) and histone acetyltransferases (Phatr3_45703 and Phatr3_45764) are highly connected with other genes, which implies that H3K4 methylation and histone acetylation may be involved in regulating genes in response to elevated *p*CO_2_. For example, the putative histone acetytransferase Phatr3_45073 interacts with genes involved in TCA cycle and carbon assimilation (Phatr3_53935, Phatr3_22122).

**Figure 3.**
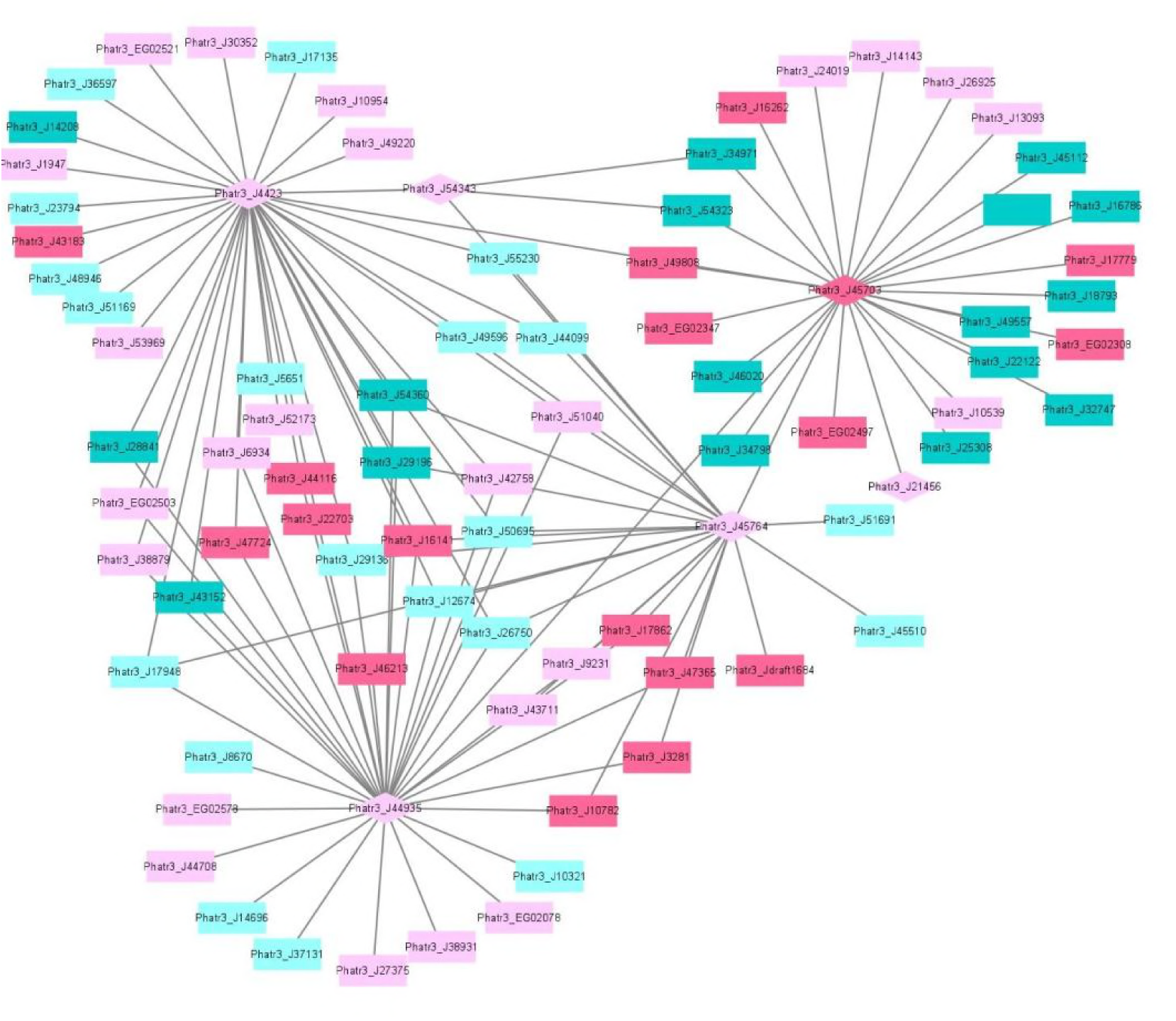
The network of differentially expressed histone modification genes and their correlated differentially expressed mRNAs under the HC and the LC conditions after growing for 15 generations. The diamonds illustrate histone modification genes while the rectangles illustrate other mRNAs. The up-regulated histone modification genes and mRNAs are filled with pink color and the down-regulated histone modification genes and mRNAs are filled with green color (UGs: dark pink, DGs: dark green, RUGs: light pink, RDGs: light green).

### The response of transcription factors to elevated *p*CO_2_

The TFs of diatoms are able to change their activity and ultimately control the diatom transcriptomic response to different signals. We investigated the expression profile of 289 genes encoding putative TFs (13, 47). Among them, 31 TFs (9 UGs, 12 RUGs, 3 DGs and 7 RDGs) were differentially expressed after growing for 15 generations in HC compared to LC (Supplementary Table 7, Figure 2). Phatr3_J37556 encoding transcription initiation factor TFIIB was the most up-regulated TF while Phatr3_J45112 encoding a heat shock factor was the most down-regulated TF. A total of seven up-regulated TFs belonged to Zn-finger and more than 50% up-regulated TFs were C2H2-type. In addition, three up-regulated TFs belonged to “Myeloblastosis” family, which may mediate signal transduction pathway in response to abiotic drivers (48). Meanwhile, 50% of the down-regulated TFs belong to heat shock factors (HSFs) which can bind to the conserved heat shock elements (HSEs) found in the promoters of target genes, including heat shock proteins (HSPs) (49). The down-regulation of HSFs correlated with the decreased expression of HSPs, an important group of molecular chaperones involved in protein assembly and folding, suggesting an acquired acclimated state under elevated *p*CO_2_.

### Long noncoding transcripts specifically induced by elevated *p*CO_2_

We analyzed the *P. tricornutum* noncoding transcriptome and identified 189 lncRNAs which are all intergenic lncRNAs (lincRNAs). LincRNAs identified in this study were compared with protein coding mRNAs generated from ssRNA-seq data for the culture growing for 15 generations under the HC and LC conditions based on Phatr3 annotation. The majority of the lincRNAs identified are between 600 and 700 nt in length and are significantly shorter than mRNAs, which range from 100 nt to more than 3000 nt. The majority of the detected *P. tricornutum* lincRNAs were also intronless, with one exon of similar size to the exons found in mRNAs previously described (13) (Supplementary Figure 2). This is a common feature of lincRNAs and has been suggested to be related to the nuclear localization of these transcripts (50). The expression levels of lincRNAs were lower than mRNA in *P. tricornutum* as shown in Supplementary Figure 3 which is also a common feature of this class of transcripts. Among them, 54 lincRNAs were up-regulated under HC while 59 lincRNAs were down-regulated. Twenty-eight up-regulated lincRNAs and 35 down-regualted lincRNAs are correlated with lincRNAs identified in the phosphate fluctuation study (13) (Figure 4). The most significantly up-regulated lincRNA was lnc_117 while the most down regulated lincRNA was lnc_145 (Supplementary Table 8).

**Figure 4.**
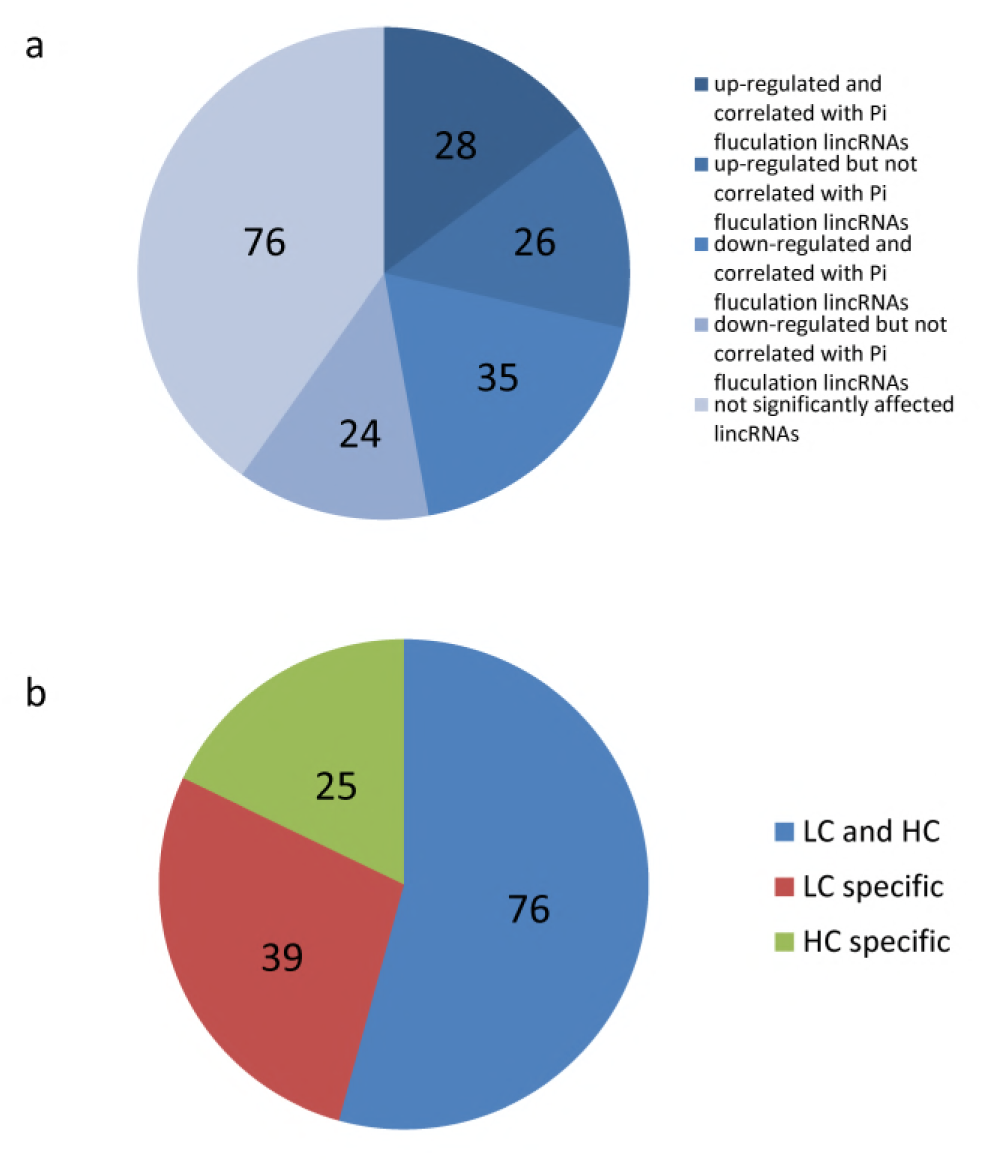
(a) The classification of lincRNAs identified in this study based on the correlation with lincRNAs identified in phosphate fluctuation study and expression. (b) The classification of lincRNAs identified in phosphate fluctuation study based on the correlation with elevated CO_2_ responsive lincRNAs.

**Figure 5.**
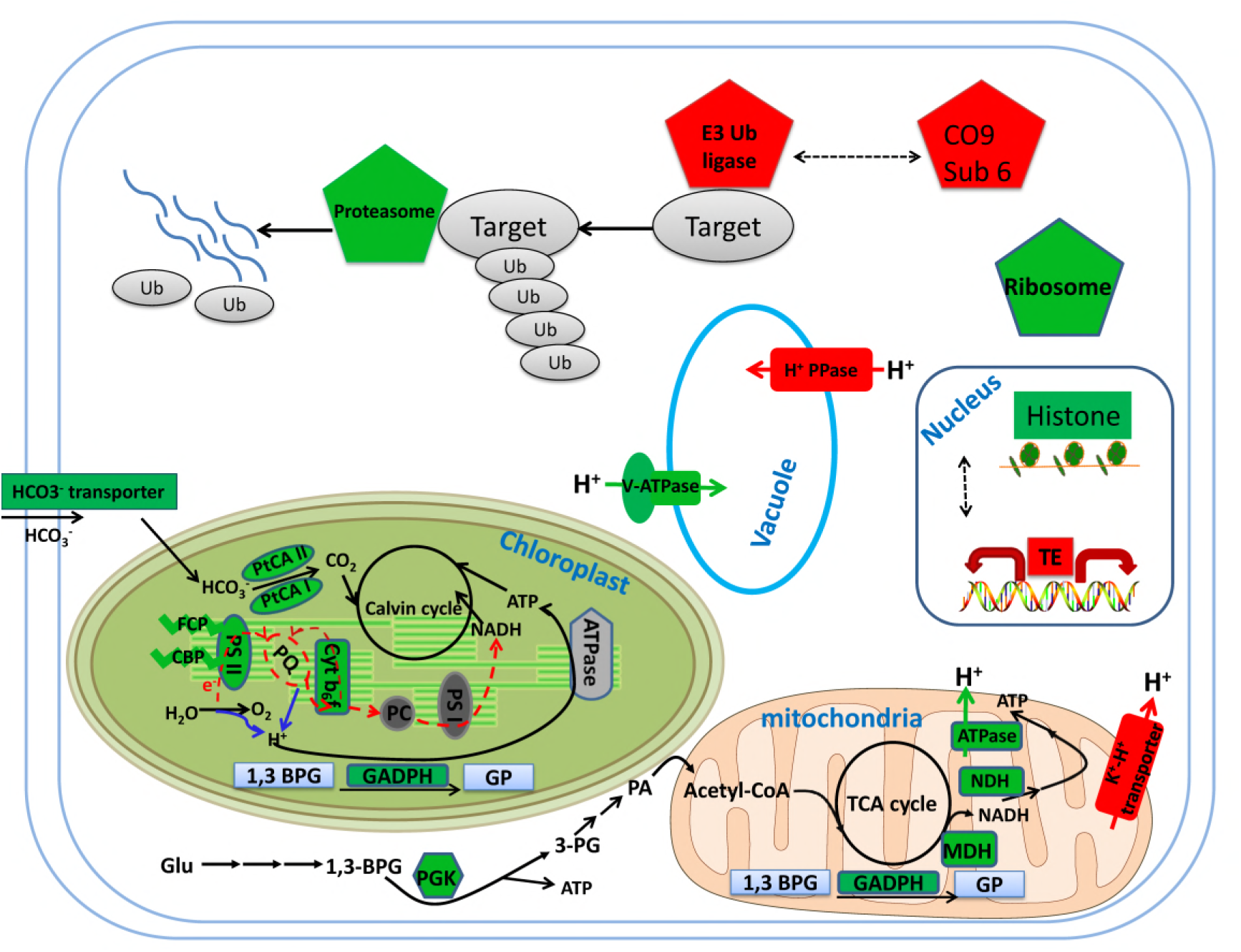
Model of metabolic and cell signaling pathways in *P. tricornutum* after acclimation to HC for 15 generations. Gene expression changes (HC/LC) are indicated by different colors: red (up-regulation) and green (down-regulation). Solid arrows indicate reactions and dashed arrows indicated regulatory relationships.

We compared the lincRNAs identified in our study with the lincRNAs associated with a phosphate fluctuation response (13). The results showed that 140 lincRNAs associated with phosphate fluctuations are also responsive to CO_2_ concentrations. Among these 140 lincRNAs, 25 lincRNAs are associated HC conditions, 39 lincRNAs are associated with LC conditions, and 76 lincRNAs are associated with both HC and LC conditions as shown in Supplementary Table 9. Most of the lincRNAs identified in this study are also involved in phosphate stress responses suggesting that linRNAs may play central regulatory roles in response to different environmental conditions. LincRNAs have previously been described to be involved in a multitude of regulatory processes at the transcriptional, post-transcriptional and epigenetic levels (12, 50, 51). LincRNAs have been namely reported to associate with several regulatory proteins such as TF and chromatin modifying complexes and have central roles in cellular homeostasis and stress response adaptations (12). Although the regulatory roles of lincRNAs in *P. tricornutum* in response to environmental stimuli remain to be determined, their expression patterns under HC and phosphate fluctuations indicate a tight regulation, which strongly suggests function.

### Conclusion and perspectives

After acclimation to the HC conditions for 15 generations, the underlying transcriptomic changes may help to maintain the physiological performance in response to elevated CO_2_. Our results suggest that chromatin-based processes may be important for the response of *P. tricorntum* to ocean acidification, not only in short term acclimation but also in long term adaptation. The striking activation of TEs combined with down regulation of histone genes indicates the potential epigenetic component in the response to ocean acidification in *P. tricornutum.* This indicates that ocean acidification may profoundly influence the adaptive processes of diatoms, important primary producers in the oceanic environment with important impacts in marine ecosystems. It was proposed that when organisms are challenged by new conditions, they can improve their fitness by a multitude of processes involving physiological acclimation, epigenetic changes, structural re-arrangements of the genome, and changes in DNA sequence, described as the adaptation spectrum. Different adaptations are characterized by the time needed for organisms to attain them and by their duration (52). Physiological adaptation is what most studies focus on, but while they may confer selective advantages, they are not actively amplified, memorized, or propagated over many generations. On the next level are epigenetic adaptations and DNA copy-number adaptations, which constitute a molecular “memory” of relatively labile genetic changes, although they do not involve changes in the actual nucleotide sequence of the genome. The observed transcriptional activation of TEs and the down-regulation of histone genes may induce further changes that influence the adaptation of *P. tricornutum* in response to elevated *p*CO_2_. In contrast with physiological adjustments, this level of adaptation could be transmitted to subsequent generations and thus have more profound effects on the whole community. It has been shown that reducing the amount of epigenetic variation available to populations can reduce adaptation in the green alga *Chlamydomonas reinhardtii*, which highlights the importance of epigenetic changes in the adaptation response to environmental changes (53).

In order to better understand the effects of ocean acidification, in addition to the ecological responses, the evolutionary responses and the underlying molecular mechanisms should be taken into account. It will be very interesting and worthwhile to investigate how epigenetic regulation and variation contribute to the acclimation in short term and adaptation in long term in response to elevated *p*CO_2_ using epigenetic approaches, such as genome wide methylome analysis and histone modification analysis (such as histone acetylation) by ChIP-seq. Furthermore, the expression profiles of several lincRNAs also suggests their involvement in the acclimation to elevated *p*CO_2_ responses which is worth to be further investigated in the future.

## Acknowledgements

This study was supported by the National Key Research and Development Program of China (grant no. 2016YFA0601302) and the National Natural Science Foundation of China (grant no. 41306096). We thank professor Senjie Lin for revising the manuscript.

## Author contribution

X.L., R. H. and K.G. planned and designed the research. R.H. performed experiments. R.H., J.D. and X. L. analyzed data. X. L., R. H., M. C. D., C. B. and L. T. wrote manuscript.

**HCO_3_^-^ transporter**: Phatr3_J54405

**Fucoxanthin-chlorophyll a-c binding protein (FCP):** Phatr3_J24119, Phatr3_J25893, Phatr3_J18049

**Fucoxanthin chlorophyll a/c protein (CBP):** Phatr3_J27278, Phatr3_J54027

**PS II**: Phar3_55057, Phatr3_J44899

**PtCA II**: Phar3_J45443

**PtCA I**: Phar3_J51305

**Cyt b6f iron-sulfur subunit**: Phatr3_J13358

**NADH dehydrogenase (NDH)**: Phatr3_EG00870,Phatr3_EG01423

**Malate dehydrogenase, mitochondrial (MDH)**: Phar3_42398

**Mitochondrial ATPase**: Phar3_J39539, Phar3_J39529

**ATPase**: Phatr3_J31133

**1,3 BPG**: 1,3 bisphosphoglycerate

**GP**:Glyceraldehyde 3-phosphate

**Glyceraldehyde 3-phosphate dehydrogenase (GADPH):** Phar3_25308 (mitochondria), Phar3_22122 (chloroplast), Phar3_32747

**Glu**: Glucose

**PG**: Phosphoglyceric acid

**PA**: Pyroracemic acid

**Phosphoglycerate kinase (PGK)**: Phar3_J48983

**Histones**: Phatr3_J34971, Phatr3_J54360, Phatr3_J46020, Phatr3_J11841, Phatr3_J26896, Phatr3_J26802, Phatr3_J34798, Phatr3_J50872, Phatr3_J11823, Phatr3_EG02092, Phatr3_EG01358, Phatr3_J50695 **Ribosome**:Phatr3_17519, Phatr3_Jdraft477, Phatr3_J28562, Phatr3_J10196, Phatr3_J47804, Phatr3_J36226, Phatr3_J51066

**Proteasome**:Phatr3_Jdraft611, Phatr3_Jdraft866, Phatr3_EG00973, Phatr3_J30003, Phatr3_J5685, Phatr3_EG02026, Phatr3_J27508, Phatr3_J49897, Phatr3_J51691, Phatr3_J20007, Phatr3_EG02638, Phatr3_J24474

